## Supplementary Figures for "SARS-CoV-2 sensing by RIG-I and MDA5 links epithelial infection to macrophage inflammation"

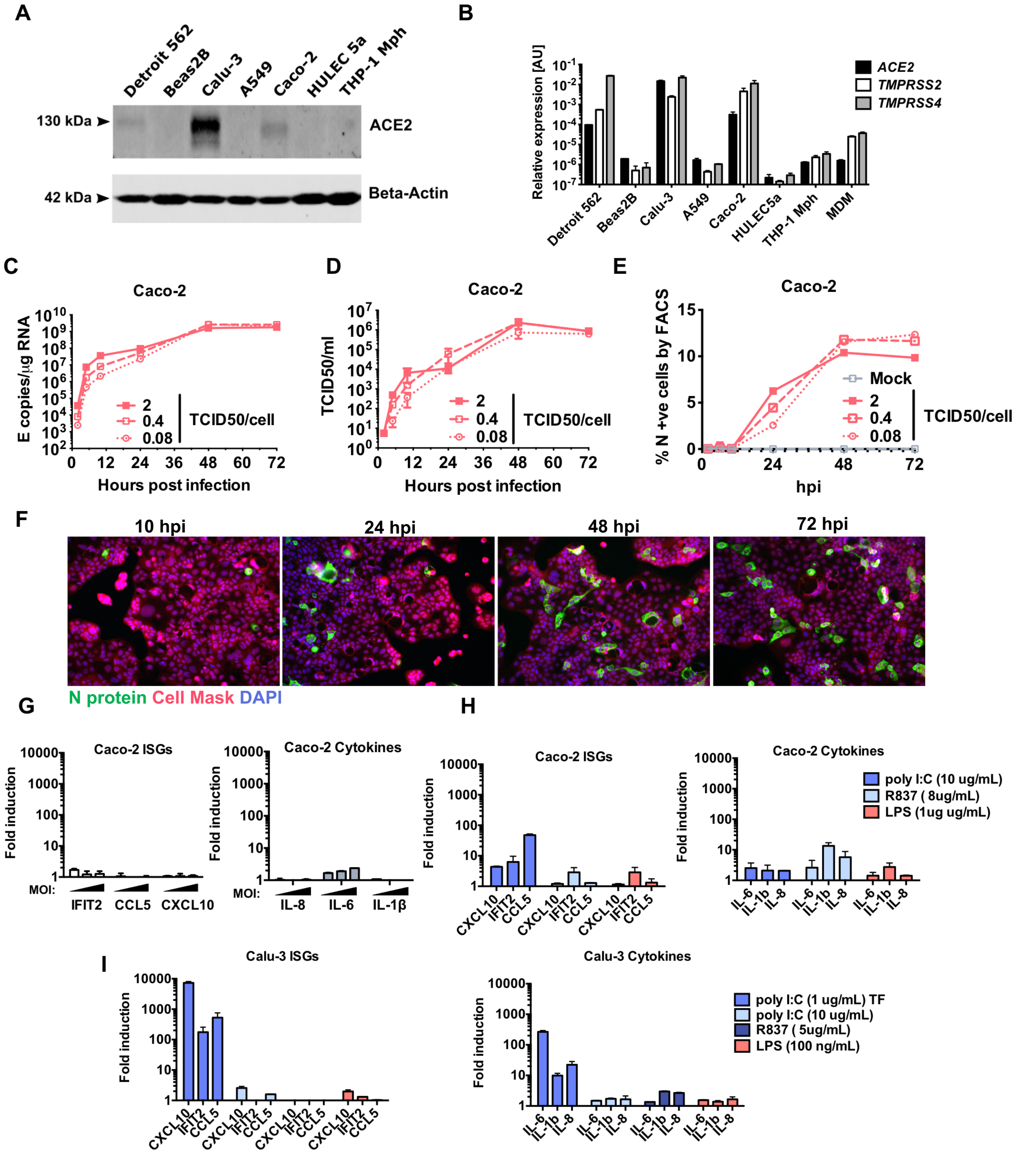


**Figure S1.** (**A**) Immunoblot detecting ACE2 expression in epithelial (Detroit 562, Beas2B, Calu-3, Caco-2), endothelial (HULEC5a) and PMA-differentiated THP-1 cells. b-Actin is detected as loading control. (**B**) *ACE2*, *TMPRSS2* and *TMPRSS4* gene expression in cell lines and primary monocyte-derived macrophages (MDM). Relative expression normalised to *GAPDH* Mean +/- SEM n=2. (**C-G**) Measurements of replication and innate immune induction in Caco-2 intestinal epithelial cells infected with SARS-CoV-2 at MOI 0.08, 0.4 or 2 $TCID50_{VERO}$/cell. Mean +/- SEM, n=2. (**C**) SARS-CoV-2 genomic and subgenomic E RNAs (qRT-PCR). (**D**) Infectious virus released from cells in (C) determined by TCID50 on Vero.E6 cells, Mean +/-SEM n=2. (**E**) Quantification of N staining from cells in (C) by flow cytometry. Mean percentage of N-positive of all live-gated cells +/- SEM, n=2. (**F**) Representative example of immunofluorescence staining of N protein (green) after SARS-CoV-2 infection of Caco-2 at MOI 0.4 $TCID50_{VERO}$/cell, at time points shown. Nuclei (DAPI, blue), cell mask (red). (**G**) Fold induction of interferon stimulated genes (ISG) of infections in (C) *IFIT2*, *CCL5*, *CXCL10* and cytokines *IL8*, *IL6*, *IL1B* at 72 hpi at MOIs $TCID50_{VERO}$/cell 0.08, 0.4 or 2, n=2. (**H**) Fold induction of ISG and cytokine gene expression in Caco-2 cells in response to innate immune activation with polyI:C, R837 and LPS for 24 h, n=2. (**I**) Fold induction of ISG and cytokine gene expression in Calu-3 cells in response to innate immune activation with polyI:C (+/- transfection, TF), R837 and LPS for 24 h, n=2. Mean +/- SEM.

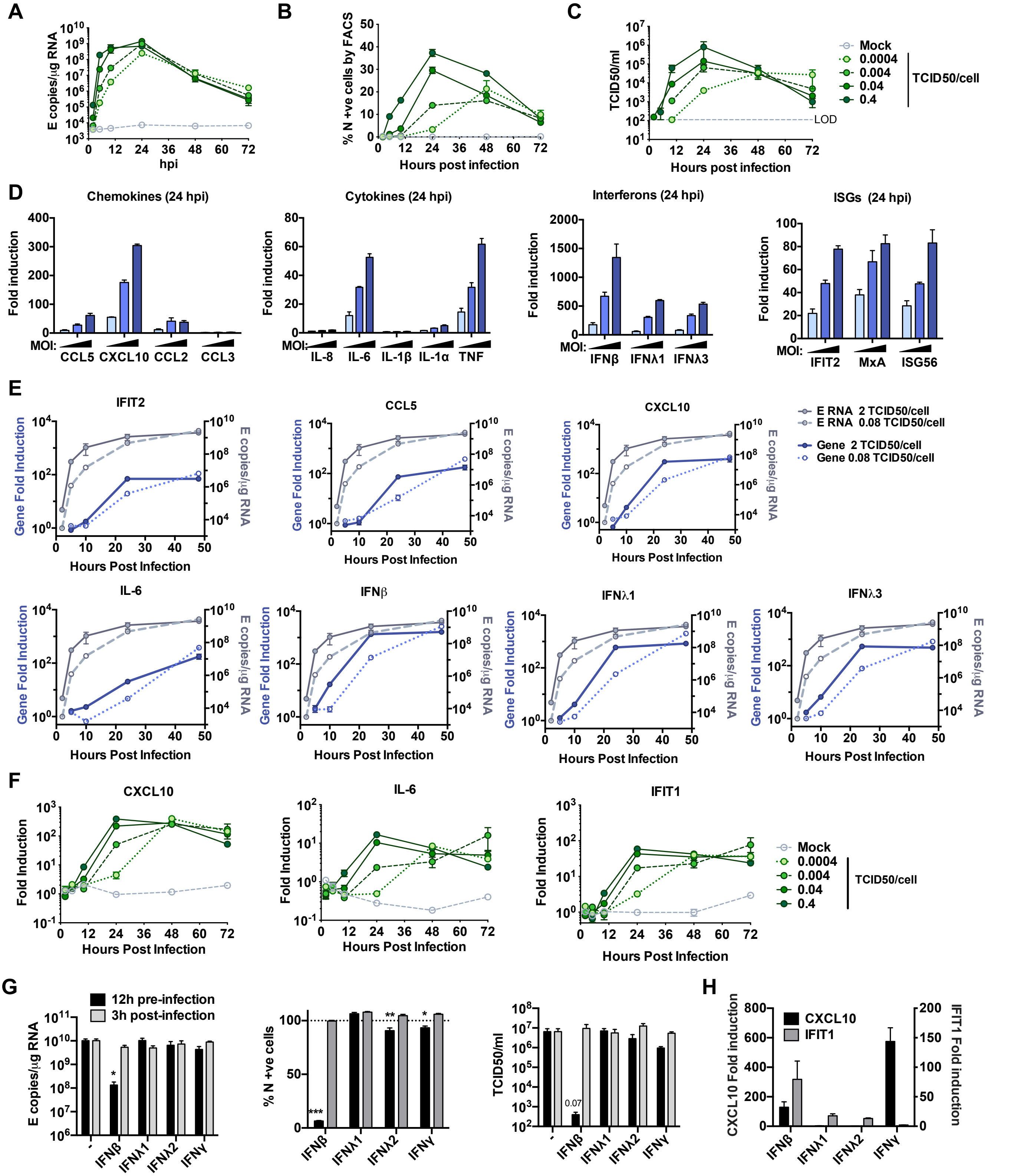


**Figure S2.**(**A-C**) Measurements of viral replication in Calu-3 lung epithelial cells infected with SARS-CoV-2 at MOIs 0.0004, 0.004, 0.04 or 0.4 $TCID50_{VERO}$/cell, n=3. (**A**) Replication of SARS-CoV-2 genomic and subgenomic E RNAs (qRT-PCR). (**B**) Quantification of N protein-positive cells from (A) by flow cytometry. Mean percentage of N +ve of all live-gated cells. (**C**) Infectious virus released from cells in (A) determined by TCID50 on Vero.E6 cells. (**D**) Fold induction of Chemokines from infections in (Figure 1) (*CCL5*, *CXCL10*, *CCL2*, *CCL3*), Cytokines (*IL-8*, *IL-6*, *IL-1*β, *IL1*α, *TNF*), Interferons (*IFN*β, *IFN*λ*1*, *IFN*λ*3*) and ISGs (*IFIT2*, *MX1*, *IFIT1*) at 24 hpi in Calu-3 cells infected at MOIs 0.08, 0.4 or 2 $TCID50_{VERO}$/cell, n=2. (**E**) Fold induction of *IFIT2*, *CCL5*, *CXCL10*, *IL6*, *IFN*β, *IFN*λ*1*, *IFN*λ*3* in Calu-3 cells at MOI 0.08 or 2 $TCID50_{VERO}$/cell each overlaid with SARS-CoV-2 E (qRT-PCR), n=2. (**F**) Fold induction of *CXCL10*, *IL-6* and *IFIT1* in SARS-CoV-2 infected Calu-3 cells from (A) at MOIs 0.0004, 0.004, 0.04 or 0.4 $TCID50_{VERO}$/cell, n=3. (**G**) SARS-CoV-2 infection (MOIs 0.04 $TCID50_{VERO}$/cell) in Calu-3 cells after addition of 10 ng/ml IFNβ, IFNλ1, IFNλ2 or IFNγ before or after infection at time points shown, measured by E RNA copies, N-positive cells (relative to untreated infection) and released virus as $TCID50_{VERO}$/ml, all measured at 24 hpi. Treatments were compared to untreated SARS-CoV-2 infected Calu-3 cells by T test. *, p<0.05; **, p<0.01; ***, p<0.001 or exact p-value are shown. Mean +/- SEM shown, n=3. (**H**) Fold induction of *CXCL10* and *IFIT1* in interferon-treated Calu-3 cells at 24h. Means +/- SEM, n=3.

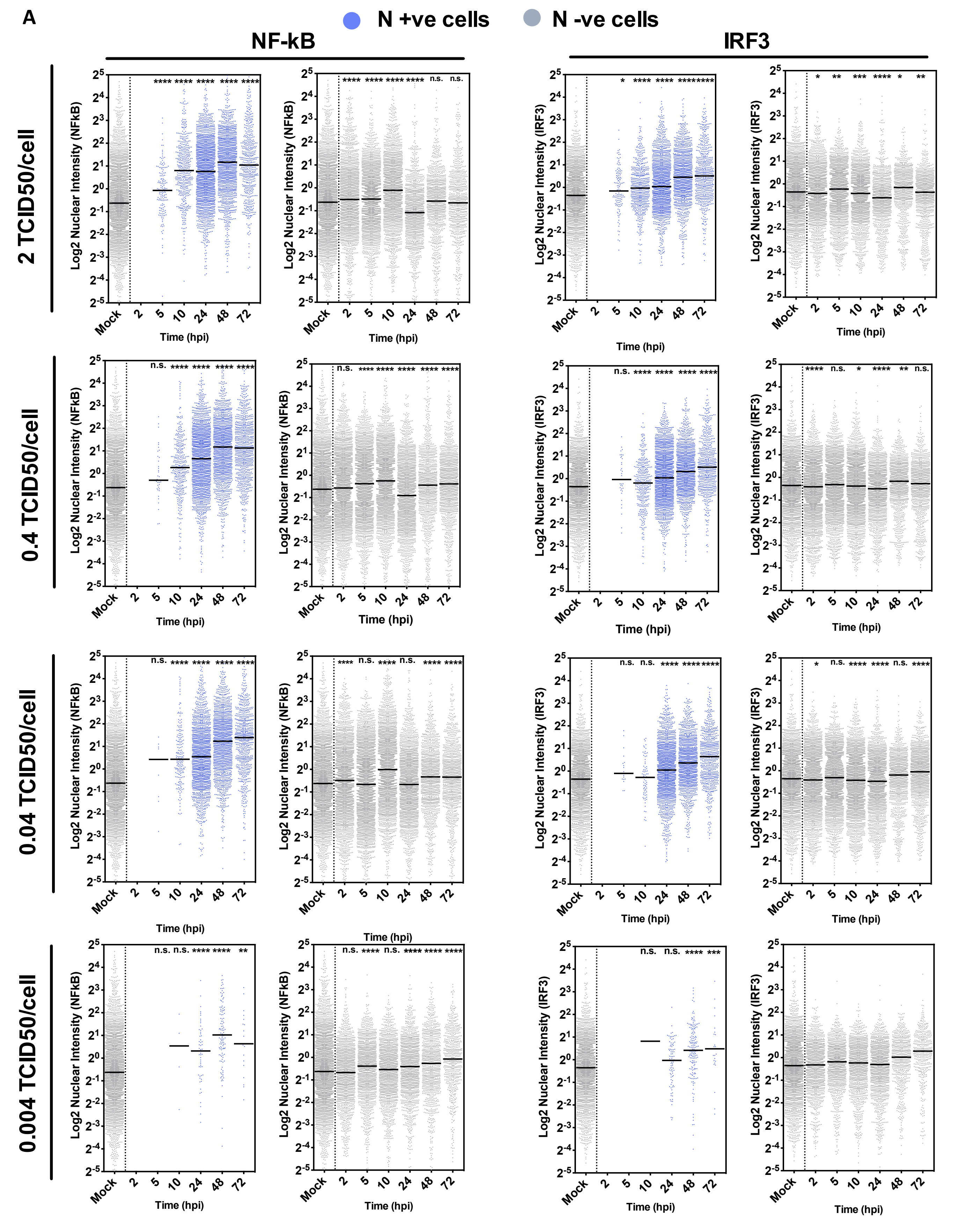


**Figure S3. (A)** Single cell analysis time course quantifying the Integrated Nuclear Intensity of NF-kB p65 or IRF3 in SARS-CoV-2 infected Calu-3 cells at MOI 2, 0.4 or 0.04 $TCID50_{VERO}$/cell as labelled. At all timepoints, nuclear intensities of NF-kB or IRF3 in nucleocapsid protein-positive infected cells (blue) and N-ve cells (grey) are shown. Nuclear Intensities of uninfected cells (Mock) at 24 h are shown as comparator. Horizontal lines indicate the mean. Kruskal-Wallis test with Dunn's multiple comparison, *, p<0.05; **, p<0.01; ***, p<0.001.

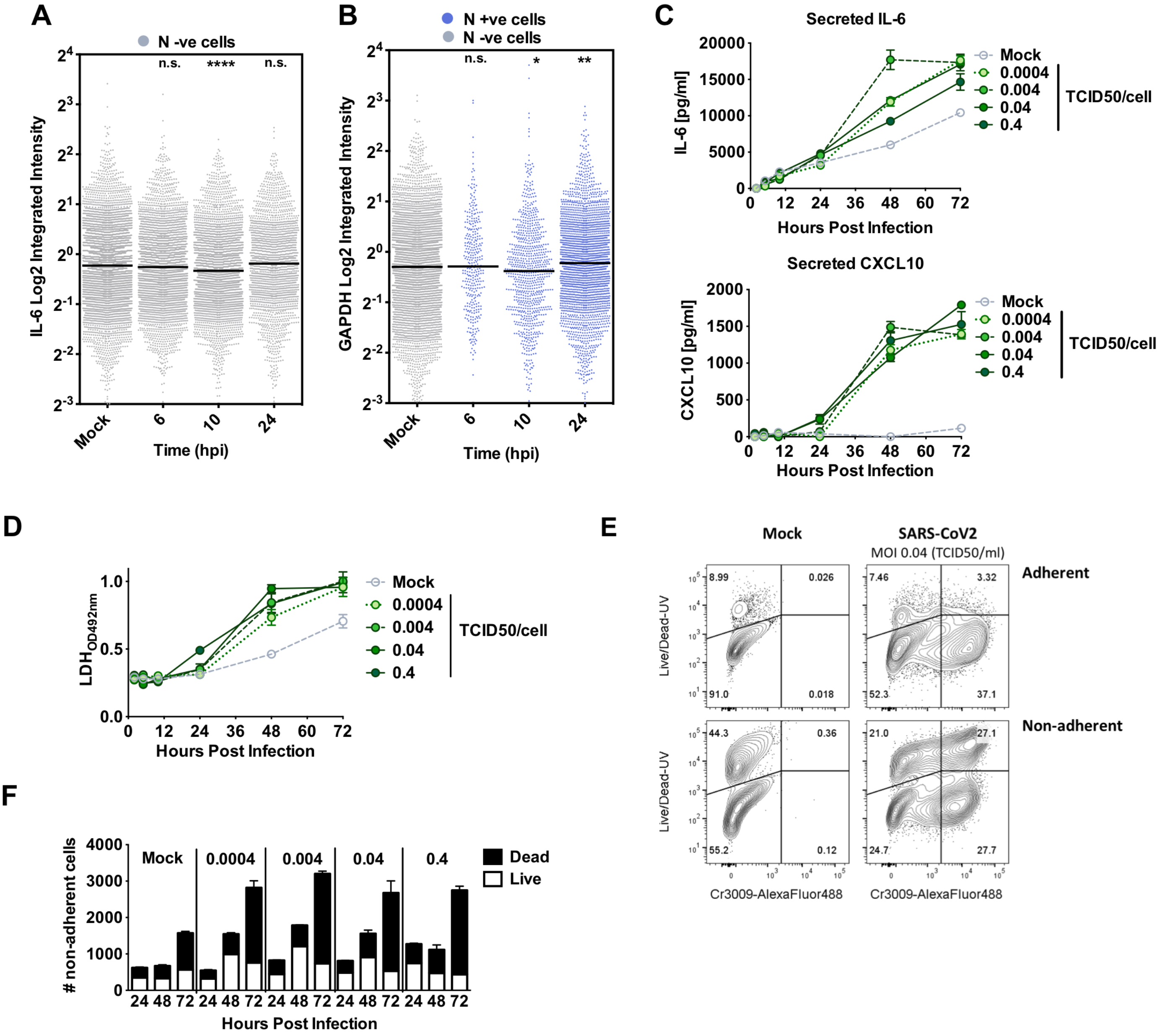


**Figure S4.** (**A**) Representative single cell RNA FISH analysis time course quantifying the Integrated Intensity of *IL-6* in uninfected (Mock) or uninfected bystander cells (uninfected cells, grey) of Calu-3 cells infected at MOI 0.4 TCID50/cell. (**B**) Representative single cell RNA FISH analysis time course quantifying the Integrated Intensity of *GAPDH* in uninfected (Mock), nucleocapsid protein-positive infected (blue) and uninfected bystander (grey) Calu-3 cells at MOI 0.4 TCID50/cell. (**A,B**) Horizontal lines indicate the median with Kruskal-Wallis test with Dunn's multiple comparison, *, p<0.05; **, p<0.01; ***, p<0.001. (**C**) Secretion of IL-6 and CXCL10 (ELISA) by infected Calu-3 cells (MOIs 0.0004, 0.004, 0.04 and 0.4 TCID50$_{VERO}$/cell), matching infections in Figure S2A-C and F. Mean +/- SEM, n=3. (**D**) Lactate dehydrogenase (LDH) release into culture supernatants by mock and SARS-CoV-2 infected Calu-3 cells (MOIs 0.0004, 0.004, 0.04 and 0.4 TCID50$_{VERO}$/cell, matching infections in Figure S2A-C and F) quantified by absorbance (492nm), means +/- SEM, n=3, (**E**) Representative flowcytometry contour plots depicting intracellular nucleocapsid protein detection (Cr3009-AlexaFluor488) and Live/Dead (Live/Dead-UV) staining. Shown are infected (MOI 0.04 TCID50$_{VERO}$/cell) and uninfected (Mock) Calu-3 cells at 48h post infection. Adherent and non-adherent cells were collected and acquired. (**F**) Quantification of Live/Dead staining of non-adherent cells recovered from supernatants of Mock or SARS-CoV-2 infected Calu-3 cultures (MOIs 0.0004, 0.004, 0.04 and 0.4 TCID50$_{VERO}$/cell) at 24, 48 or 72 hpi. Mean +/- SEM, n=3.

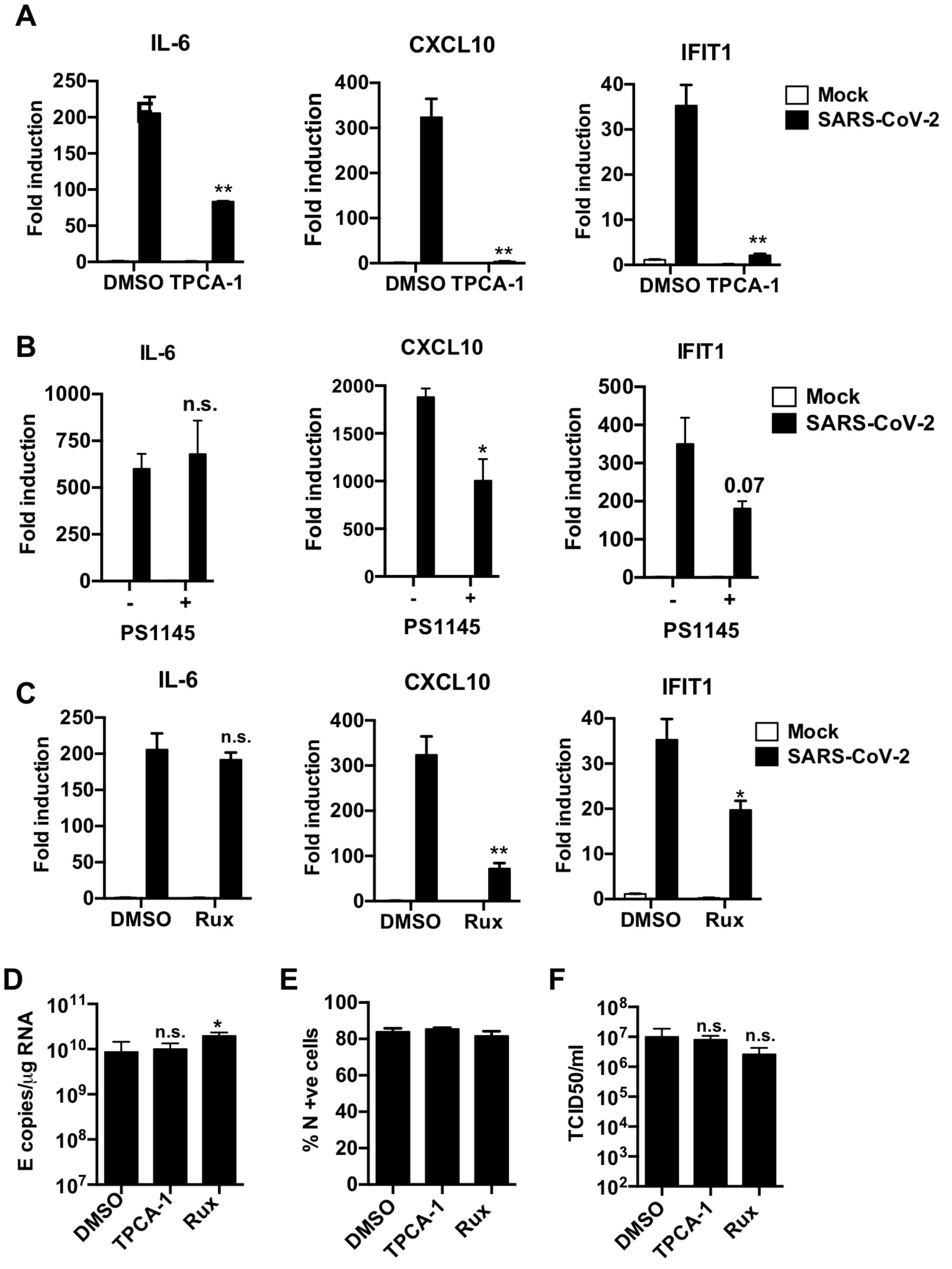


**Figure S5. (A-C)** Fold gene induction of *IL-6, CXCL10* and *IFIT1* 24hpi of Calu-3 with SARS-CoV-2 (MOI 0.04 $TCID50_{VERO}$/celll) infected in the presence of: **(A)** 10 µM TPCA-1 (**B**) 10 µM PS1145 or (**C**) 10 µM Ruxolitinib (Rux) with DMSO as control in each case. (**D-F**) Measurement of SARS-CoV-2 (MOI 0.04 $TCID50_{VERO}$/cell) replication in Calu-3 in the presence or absence of 10 µM TPCA-1, 10 µM Ruxolitinib (Rux) or DMSO vehicle as shown measuring (**D**) genomic and subgenomic E RNA, (**E**) N +ve cells by flow cytometry, (**F**) released virus in supernatant ($TCID50_{VERO}$/cell) at 24 hpi. Mock and SARS-CoV-2 infected/treated conditions were compared by two tailed t test. *, p<0.05; **, p<0.01; n.s., non-significant. Mean +/- SEM, n=3.

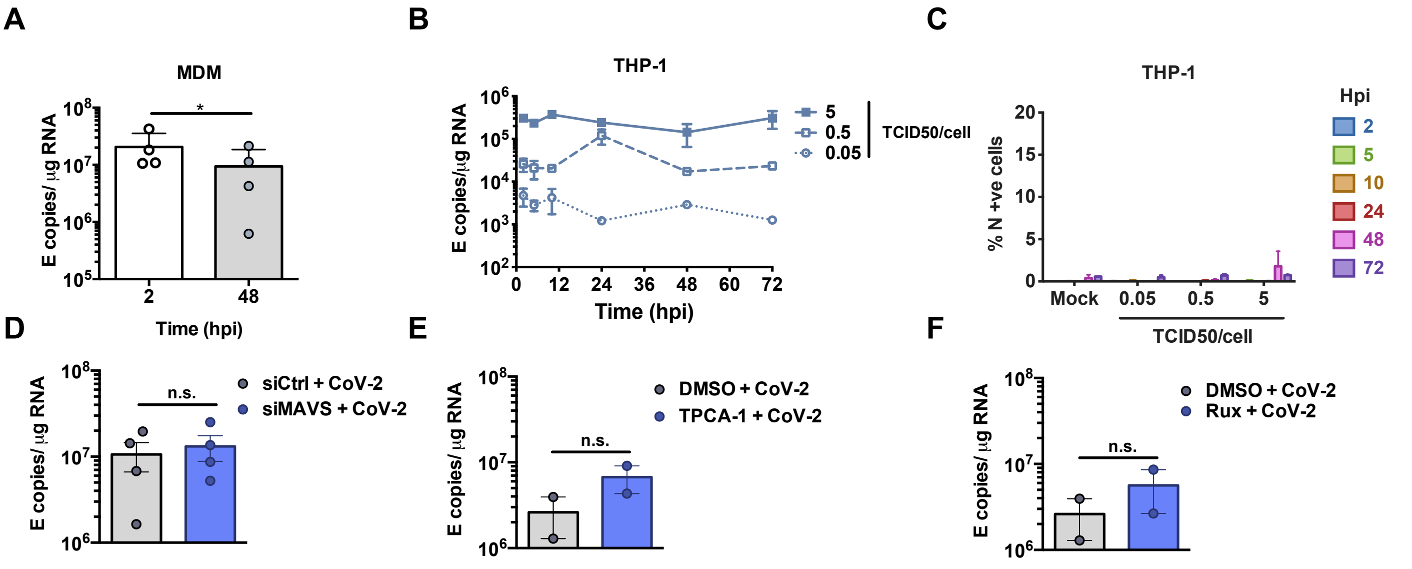


**Figure S6.** (**A**) MDM were exposed to SARS-CoV-2 for 2 or 48h and viral genomic and subgenomic E RNA measured (RT-qPCR), means shown +/- SEM, n=4. Statistical comparison between 2 and 48h by paired two-tailed t test, *, p<0.05. (**B, C**) PMA-differentiated THP-1 macrophages were exposed to SARS-CoV-2 for 72h measuring (**B**) viral genomic and subgenomic E RNA (qRT-PCR) or (**C**) N protein +ve cells (flowcytometry). Mean+/- SEM, n=2. (**D-F**) Detection of viral genomic and subgenomic E RNA extracted from MDM exposed for 48 h to SARS-CoV-2-containing conditioned medium from Calu-3 cells infected in the presence or absence of (D) MAVS depletion by siRNA, (E) 10 µM TPCA-1 or (F) 2 µM Ruxolitinib. Mean +/- SEM n=2-4. Statistical comparison by paired two-tailed t test, n.s. : non significant.

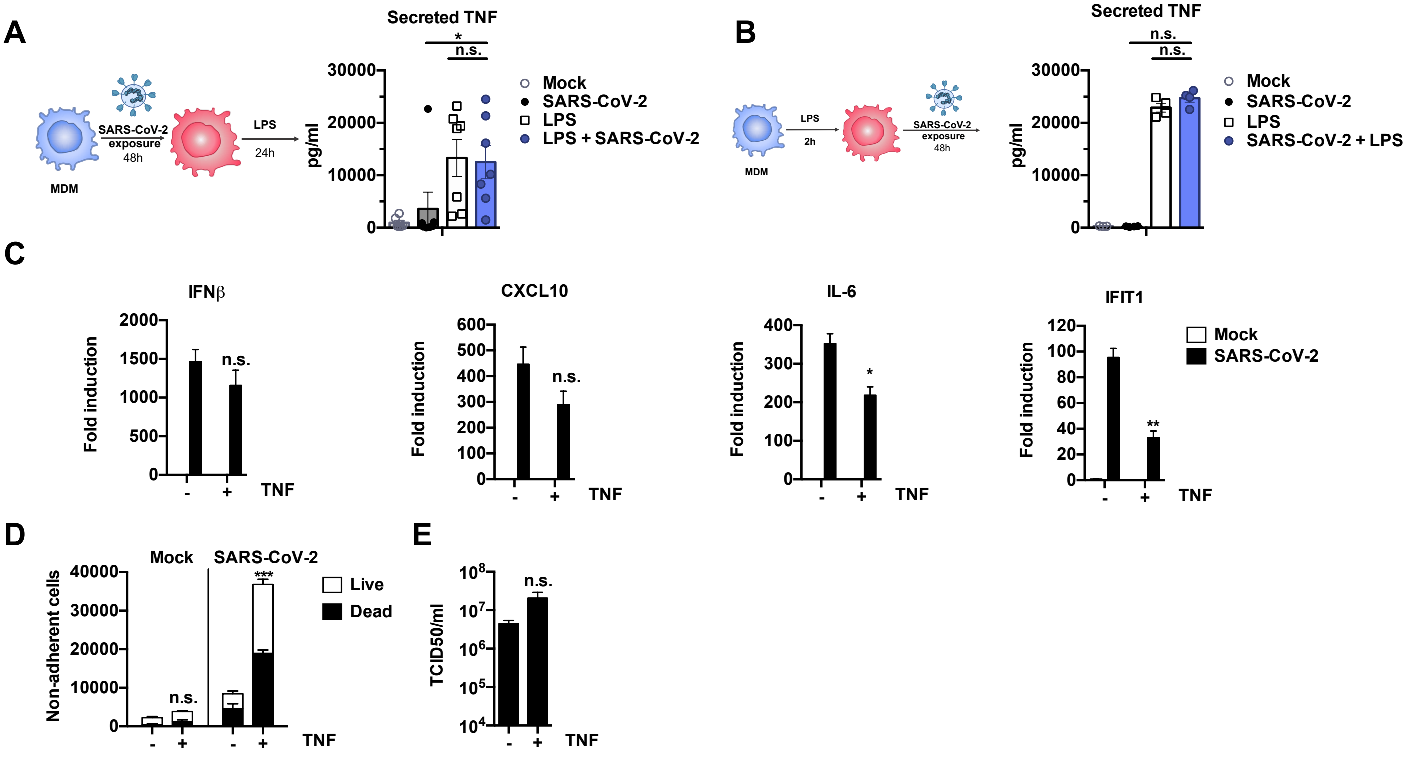


**Figure S7.** (**A-B**) Schematic of experimental design showing timing and order of exposure of MDM to LPS and SARS-CoV-2 (as in figure 6), and measurement of TNF (ELISA). Groups were compared as indicated by Wilcoxon matched-pairs signed rank test, *, p<0.05. Mean+/-SEM, n=4-7. (**C-E**) SARS-CoV-2 infection of Calu-3 cells (MOI 0.04 $TCID50_{VERO}$/cell) in the presence or absence of 10 ng/ml TNF added at the time of infection. SARS-CoV-2 infected TNF + and – groups were compared by two tailed t test. *, p<0.05; **, p<0.01; n.s., non-significant. Mean +/- SEM shown, n=3. (**C**) Fold gene induction of *IFN*β, *CXCL10, IL-6,* and *IFIT1* at 24hpi. (**D**) Quantification of Live/Dead staining of non-adherent cells recovered from culture supernatants at 24hpi. (**E**) Virus titres in Calu-3 supernatants at 24 hpi ($TCID50_{VERO}$/cell).
