## Supplementary Tables for "SARS-CoV-2 sensing by RIG-I and MDA5 links epithelial infection to macrophage inflammation"

| Supplementary table 1A. Analysis parameters for detection of SARS-CoV-2 infection by intracellular nucleocapsid protein in Calu-3 cells. | |
|---|---|
| **Viral infection (Intracellular Granules Module, 10X/0.4 NA)** | **Analysis Parameters:** |
| Nucleus : Nucleus Smooth | 1.9 |
| Nucleus : Nucleus Background Subtraction | 175 |
| Nucleus : Nucleus Intensity Threshold | 310 |
| Nucleus : Nucleus Max Patch Size | 100 |
| Nucleus : Nucleus Maximum Area | 800 mic. |
| Nucleus : Nucleus Minimum Area | 32 mic. |
| Cell : Cell Smooth | 5 |
| Cell : Cell Background Subtraction | 2000 |
| Cell : Cell Intensity Threshold | 5000 |
| Granules : Granules Smooth | 5 |
| Granules : Granules Background Subtraction | 45 |
| Granules : Granules Intensity Threshold | 930 |
| Granules : Granules Maximum Area | 1000 mic. |
| Granules : Granules Max Patch Size | 20 |
| Granules : Granules Minimum Area | 10 mic. |
| Population: Infected: Mean Granule Inensity | ≥1000 |
| Output | Cell Count, Nuclear Area, Nuclear Intensity (DAPI), Cell Area (CellMask), Granule Area, Granule Intensity (N-Protein) |

| Supplementary table 1B. Analysis parameters for IRF3/NFkB nuclear intensity during SARS-CoV-2 infection in Calu-3 cells. | |
|---|---|
| **IRF3/NFkB Nuclear Intensity (Intranuclear Foci Module, 40X/0.75NA)** | **Analysis Parameters:** |
| Nucleus : Nucleus Smooth | 1 |
| Nucleus : Nucleus Background Subtraction | 109.8 |
| Nucleus : Nucleus Intensity Threshold | 150 |
| Nucleus : Nucleus Maximum Merge Area | 60 |
| Nucleus : Nucleus Minimum Area | 30 |
| Nucleus : Nucleus Maximum Area | 900 |
| Foci : Foci Smooth | 1 |
| Foci : Foci Background Subtraction | 105 |
| Foci : Foci Intensity Threshold | 2170 |
| Foci : Foci Maximum Merge Area | 105 mic. |
| Foci : Foci Minimum Area | 30 mic. |
| Foci : Foci Maximum Area | 2000 mic. |
| Output: | Cell Count, Nuclear Area, Nuclear Intensity (DAPI), Nuclear Intensity of Marker Protein (IRF3/NFkB), Foci Area, Foci Intensity (N-Protein) |

| Supplementary table 1C. Analysis parameters for RNA FISH detection during SARS-CoV-2 infection in Calu-3 cells. | |
|---|---|
| **RNA FISH analysis (Mitochondria Module, 40X/0.75 NA)** | **Analysis Parameters:** |
| Nucleus : Smooth | 1.2 |
| Nucleus : Background Subtraction | 68 |
| Nucleus : Intensity Threshold | 220 |

| | |
|---|---|
| Nucleus : Maximal area for merging | 30 |
| Nucleus : Minimum Area | 57 mic. |
| Nucleus : Maximum Area | 800 mic. |
| Cell : Smooth | 2 |
| Cell : Intensity Threshold | 2500 |
| Mitochondria : Background Subtraction | 108.8 |
| Mitochondria : Minimum Length | 1 |
| Mitochondria : Typical Width | 1 |
| Mitochondria : Intensity Threshold | 2000 |
| Mitochondria : Maximal area for merging | 1 |
| Mitochondria : Minimum Area | 1 mic. |
| Mitochondria : Maximum Area | 121 mic. |
| Output: | Cell Count, Nuclear Area, Nuclear Intensity (DAPI), Cell Area (CellMask), Cellular Intensity of Marker Protein(IL6/IFIT1/GAPDH), Mitochondria Area/ Intensity (N-Protein) |
